# Diverse stepping motions of cytoplasmic dynein revealed by kinetic modeling

**DOI:** 10.1101/516179

**Authors:** Shintaroh Kubo, Tomohiro Shima, Shoji Takada

## Abstract

Cytoplasmic dynein is a two-headed molecular motor that moves to the minus end of microtubule (MT) using ATP hydrolysis free energy. By employing its two heads (motor domains), cytoplasmic dynein shows various bipedal stepping motions; the inchworm and hand-over-hand motions, as well as non-alternate steps of one head. However, the molecular basis to achieve such diverse stepping manners remains obscure. Here, we propose a kinetic model for bipedal motions of cytoplasmic dynein and performed Gillespie Monte Carlo simulations that reproduces most experimental data obtained to date. The model represents status of each motor domain as five states according to conformations, nucleotide- and MT-binding conditions of the domain. Also, the relative positions of the two domains were approximated by three discrete states. Accompanied by ATP hydrolysis cycles, the model dynein stochastically and processively moved forward in multiple steps via diverse pathways, including inchworm and hand-over-hand motions, same as experimental data. The model reproduced key experimental motility-related parameters including velocity and run-length as functions of ATP concentration and external force. Our model reveals that, in a typical inchworm motion, the leading domain moves via the ATP-dependent power-stroke of the linker coupled with a small change in the stalk angle, whereas the lagging domain moves via diffusion dragged by the leading domain. Moreover, the hand-over-hand motion in the model dynein clearly differs from that of kinesin by the usage of the power-stroke.

**Author Summary:** Cytoplasmic dynein is a two-headed molecular motor, which moves linearly and transports intra-cellar organelles along microtubules driven by ATP hydrolysis free energy. In contrast to other better-known molecular motors, such as kinesin, dynein is known to take various stepping motions including motions akin to human walking and inchworm-like motions. However, molecular mechanisms underpinning the diverse stepping motions are unclear. Here, based on recent high-resolution structure information and single-molecule motility assay data, we designed a kinetic model that explicitly include two heads, each of which makes ATP hydrolysis cycles and moves along the microtubules. Using the model, we performed Monte Carlo simulations. The simulation reproduced most of currently available experimental results. More importantly, the simulation suggested molecular mechanisms of various stepping motions. While stepping motions apparently resemble to those proposed before, once looking into details, we found the resulting mechanisms distinct from previously proposed ones in the usage of ATP and protein conformation changes coupled with stepping motions.

## Introduction

Cytoplasmic dynein (hereafter, denoted as dynein for simplicity) is a molecular motor that show bipedal motions on the microtubules (MT) to its minus end, driven by ATP hydrolysis free energy[1–3]. This motility enables dynein to play essential roles in various cellular functions including intracellular transport, positioning of organelles and cell division. As such, dynein is often compared with another MT-based molecular motor, kinesin, most of which walks to the plus end of MT[4]. Single-molecular measurements clarified that the kinesin-1 walks rather regularly via the so-called hand-over-hand manner; two motor domains alternately move from the lagging position to the leading position, akin to human walking. This motion of kinesin leads to precise 8 nm steps per one ATP hydrolysis reaction[5]. The hand-over-hand mechanism is also in harmony with the experiment that a mutant kinesin that impairs one motor domain severely slows down kinesin motility [6,7]. In contrast, dynein moves more stochastically with various step size, which ranges 4-32 nm with its representative size of 8 nm [8–11]. In addition to the hand-over-hand motions, dynein can take the so-called inchworm-like motions; walking via alternating steps with one motor domain being always ahead of the other domain, as well as non-alternating steps[10–13].

Previous structural studies delineate the ATP-dependent conformational changes in each dynein motor domain. The dynein motor domain consists of the ATPase associated with diverse cellular activities (AAA+) ring[14,15], the microtubule binding domain (MTBD), the stalk, the linker, and the tail domain (see cartoons in Fig 1A). The AAA+ ring (a large donut shape in Fig 1A) made of six subdomains, AAA1-AAA6, possesses the ATPase activity and energizes the dynein movement[16]. The MTBD (small circles near the MT in Fig 1A) is a small domain responsible for the binding to MT. The stalk is a long coiled-coil that connects the AAA+ ring and the MTBD[16,17]. The linker (an arrowhead-like object drawn on the AAA+ ring) is associated with the AAA+ ring and connects to the tail domain where two motor domains dimerize (the tail domain not drawn in Fig 1A)[18,19]. Biochemical and structural experiments have revealed that ATP hydrolysis reactions in AAA1 play the primary role in the movement, affecting conformations of the entire motor domain[15,20]. In particular, depending on the nucleotide state in AAA1, both the linker and the MTBD make marked structural changes which are of central importance in the motility[17]. When AAA1 is in the ATP-bound state, the linker is largely bent on the AAA+ ring and the tip of the linker is located near AAA2/AAA3 subdomains (the bottom-left cartoon in Fig 1B), whereas, in the ADP bound and apo state of AAA1, the linker tends to be more extended on the AAA+ ring reaching to the AAA4/AAA5 subdomains (the top-right cartoon in Fig 1B)[21–24]. The linker is expected to swing simultaneously or immediately after Pi release and swings back upon ATP binding, which are termed the power-stroke and recovery-stroke, respectively. The power-stroke of the linker is considered to be responsible for dynein force generation [17,25,26]. The MTBD tends to take a low affinity state to MT in the ATP bound state, while it takes a high affinity conformation to MT in the ADP-bound and apo states [20]. This ATP-dependent change in the affinity of MTBD must be critical, because dynein needs to bind tightly with MT when it exerts force against MT but needs to dissociate from MT during recovery-stroke to avoid backward movement[27,28]. In addition to these specific parts of the motor domain, allosteric conformational changes in the entire motor domain may play an important role for the dynein movement[29,30].

**Fig 1.**
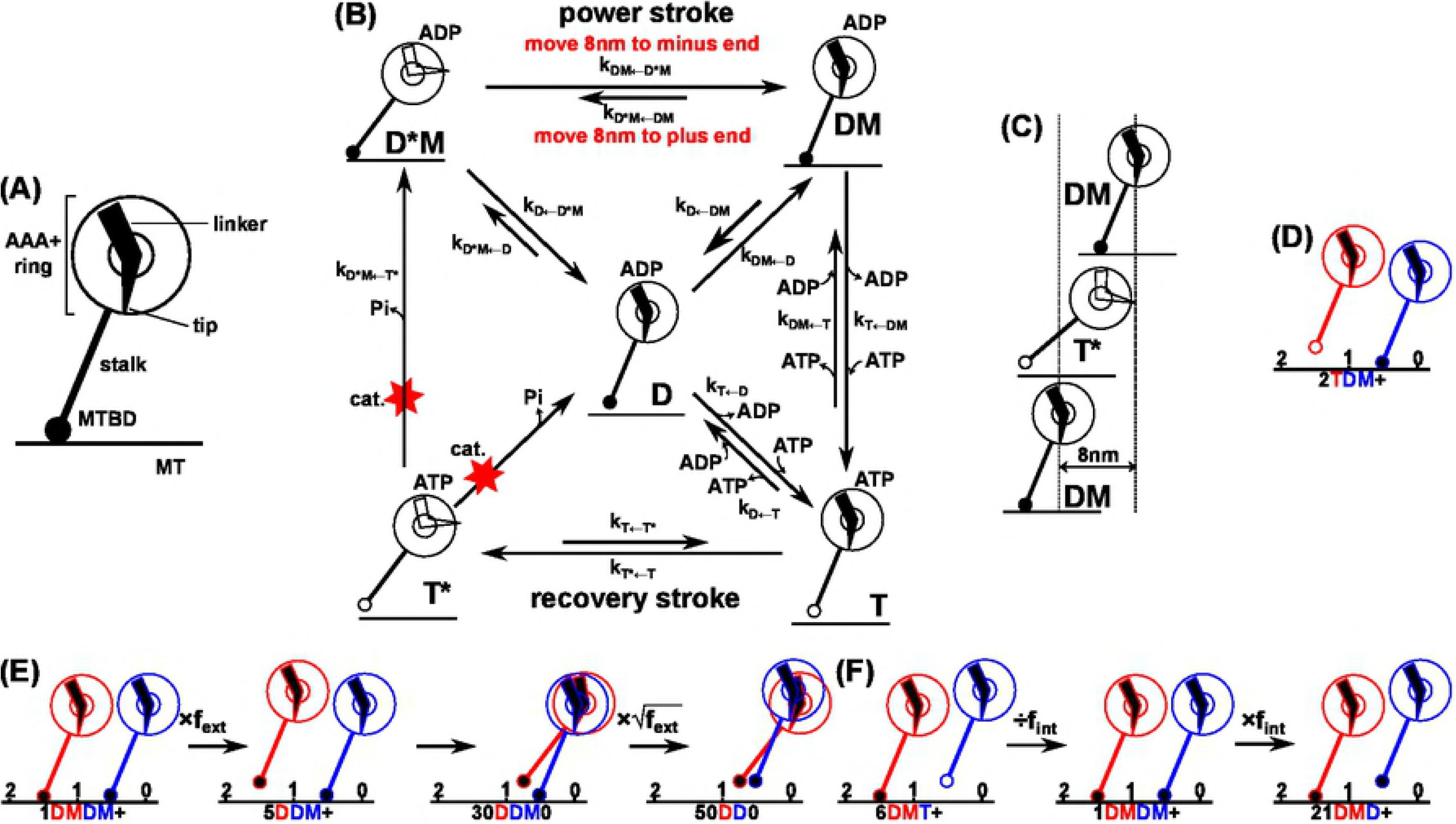
The kinetic model of the two-headed dynein. (A) Architecture of the dynein motor domain structure. (B) The five state model of one dynein motor domain and transitions among states. (C) The motor domain position. The position is defined by the tip of the linker where cargos and/or beads are attached. In the model, it takes discrete positions with 8 nm spacing. (D) The dimeric dynein state. The first (designated red) motor domain is either 8 nm ahead of, at the same position as, or 8 nm behind the second (blue) motor domain, which are represented, respectively, as +, 0, or –. As a whole, there exist 5 x 5 x 3 = 75 states for the bipedal dynein. We label each state, e.g., 2TDM+ where the first integer is the state number from 1 to 75, followed by the states of the red and then blue motor domains. The rightmost symbol represents the positioning of the red domain relative to the blue. (E) Effects of external force to the backward direction on transition rates. (F) Effects of internal force/tension on transition rates.

However, these ATP-dependent conformational changes in each motor domain alone do not explain how diverse walking manners are realized. In order to clarify the mechanism of various walking manners, we need to characterize coordination of the two motor domain movements coupled with ATPase cycle. Observing the coordinated motion directly by single-molecule experiments is, however, currently difficult due to the time- and the spatial-resolution.

Therefore, several theoretical models have been proposed to understand the walking mechanisms of dynein. Most theoretical models proposed before the X-ray structure reports are unavoidably simple and consider only one motor domain [31,32]. A more elaborate kinetic model that explicitly deals with coordination of the two motor domains clarified a class of necessary coordination and the force-dependent motility change well [33]. The model is, however, limited to the hand-over-hand coordination and is not compatible with recent experimental data. More recently, a mechano-chemical model that connects the two ATP-dependent motor domains via elastic bonds elucidated correlation between the motion and the tension [34]. Yet, in the model, chemical cycles in the two domains are treated as independent. However, in order to clarify the mechanism of versatile walking mechanisms, it is crucial to model how ATP hydrolysis reactions and conformational change proceed, correlated with the relative configuration of the two motor domains.

In this study, we propose a new kinetic model of dynein that explicitly contains chemical and conformational states of each motor domain, directly coupled with the relative positions of the two domains along MT. The kinetic model can explain molecular basis of versatile walking manners. We represent chemical and conformational states of each dynein motor domain as 5 discrete states. For the relative positions of the two domains, for simplicity, we assume it taking 3 discrete states: either one is 8 nm ahead of, at the same position as, or 8 nm behind the other motor domain along MT. Together our kinetic model consists of 5 × 5 × 3=75 states and considers transitions among these states, which is solved by Monte Carlo (MC) simulations with the Gillespie algorithm[35]. We reproduced experimentally observed behaviors with various dynein walking manner and uncovered detailed and comprehensive pathways.

## Model and Methods

### Five state model for a dynein motor domain

Our model assumes that each dynein motor domain takes one of five possible states depending on the combination of the nucleotide bound in AAA1, the binding to MT, conformation of the linker, and that of MTBD (Fig 1B). The five states are the followings:

1. The DM state: ADP is bound in AAA1. The linker is in the post-power-stroke (extended) state. The high-affinity MTBD binds to MT.
2. The T state: ATP is bound in AAA1. The linker is in the post-power-stroke state. The low-affinity MTBD is unbound from MT.
3. The T* state: ATP is bound in AAA1. The linker is in the pre-power-stroke (bent) state. The low-affinity MTBD is unbound from MT. The stalk leans towards the MT long axis.
4. The D*M state: ADP is bound in AAA1. The linker is in the pre-power-stroke state. The high-affinity MTBD binds to MT. The stalk leans towards the MT long axis.
5. The D state: ADP is bound in AAA1. The linker is in the post-power-stroke state. The high-affinity MTBD is, however, unbound from MT.

Of the five states, the states (1) and (3) correspond to X-ray crystal structures solved to date (PDB ID: 3VKH and 4RH7, respectively)[22,36] (note that those are for different families) and thus are assumed to be relatively stable, whereas the other three states are considered as transient states with higher free energies. We note that, when the linker takes the pre-power-stroke state (the D*M and T* states), we assume that the stalk leans to the MT long axis with a smaller angle between the stalk and the MT compared with the other states. This is based on experimental data that suggests the motor domain to stand up with the MTBD position fixed upon power stroke motion[37]. This suggestion was further supported by some electron-microscopic observations for cytoplasmic dynein [38,39] as well as for axonemal dynein [17].

Next, we setup transitions among the five states of motor domain.

1. DM→T: ADP release is followed by ATP binding and a dissociation of the motor domain from MT
2. T→T*: The recovery-stroke of the linker
3. T*→D*M: ATP hydrolysis is followed by the Pi release and the binding of the motor domain to MT
4. D*M→DM: The power-stroke of the linker The above four transitions form the main (and thus productive) ATP hydrolysis cycle. Additionally, we include the following off-pathways.
5. D*M→D & DM→D: The dissociation of the motor domain from MT
6. D→T: ADP release is followed by ATP binding, without the binding to MT
7. T*→D: ATP hydrolysis is followed by Pi release, without the binding to MT.

Our model assumes that the ATP hydrolysis cycle of one ATP molecule corresponds to one power-stroke and recovery stroke, without considering ATP hydrolysis in other AAAs than AAA1 subdomain.

Since the dynein motor domain never synthesize ATP from ADP and Pi in any condition, we do not consider the reverse transitions of (3) or (7), D*M→T* or D→T*. For all the other transitions, we take into considerations of the reverse transitions. We note that all the cyclic transition paths that do not include the ATP hydrolysis, i.e. T*→D*M and T*→D, should give no net free energy changes. This leads us to the following equalities to be satisfied.

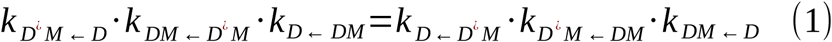

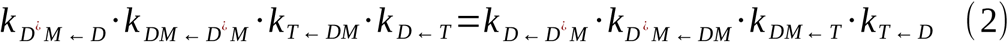

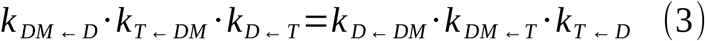

### Movement of dynein motor domain along MT

We next define the position of the dynein motor domain. Since cargos *in vivo* and beads in single-molecule experiments are connected to the tip of the linker, we define the position of motor domain by the position of the tip of the linker. In addition, we approximate that the tip of the linker resides in discrete positions spaced with 8 nm (Fig 1C).

Next, we discuss the movement of the motor domain. We assume that the position of the linker tip changes via the power-stoke/recovery-stroke, and the diffusive motion. First, in the power-stroke transition D*M→DM (see Fig 1B), the tip of the linker position moves 8 nm to the forward direction (the minus end of MT), whereas in the reverse reaction of the power-stroke (note that this is not the recovery stroke), the tip moves 8 nm to the backward direction (the plus end of MT) although the reverse reaction rarely happens.

Both the yeast and human dynein molecules can move forward against 5∼10 pN backward loading [9,40]. If the load force disturbs the linker motion completely, the dynein molecules would not step forward. Therefore, as a common feature of dynein, the power-stroke motion of the linker should be able to undergo against more than 5∼10 pN of the load. This led us to model the following energy difference between pre- and post-power stroke states;

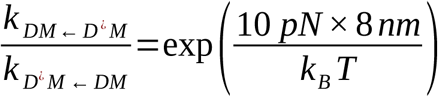

where *T* is the temperature, the Boltzmann constant *k* _*B*_ takes *k* _*B*_=0.0138 *pN ·nm* / *K.* We note that the tip of the linker does not move during the recovery-stroke from the T to T* states because the MTBD is unbound from MT.

Next, we consider the diffusive motion. In our five states model, only in the D, T and T* states among five states of our model, the motor domain can move diffusively along MT. We assume that the motor domain takes an 8-nm step to forward or backward direction during the diffusive movement. We define the rates for the forward and backward diffusive transitions as *λ*_*for*_ and *λ*_*back*_, respectively. The values of *λ*_*for*_ and *λ*_*back*_ are set as the same for all the three states; the D state, the T state, and the T* state. In addition, by symmetry, *λ*_*for*_ and *λ*_*back*_ must be the same values when the external force is not applied.

### The dimeric dynein motor domains

We assume that dynein moves as a homo-dimer of the motor domain throughout this study. To distinguish two identical motor domains, throughout the paper, we call the first one as “red domain” and the second domain as “blue domain”, merely for convenience, corresponding to the colors used in figures (Fig 1D for example). For the minimal complexity of the model, we set that the difference in the positions of the two motor domains along MT being either +8 nm, 0 nm, or −8 nm (Fig 1D for the case of +8 nm). Thus, the relative positions of the two dynein motors can be regarded as the three-state model. When the red motor domain is ahead of, at the same position as, and behind the blue motor domain, we write their configurations as +, 0, and -, respectively (The minus end of MT is regarded as the forward direction). Assumed here is that the two motor domains use different protofilaments so that they can have 0 nm distance along axial direction of MT. We do not include the side-steps from one to another proto-filaments. We also note that the generalization of this restraint of the inter-motor-domain distance to the multiples of 8 nm is straightforward.

Combining 5 states for each motor domain and 3 states for the relative positions between the two motor domains, the current model for the dimeric dynein has 5 ×5 ×3=75 states in total. To indicate each of 75 states clearly, we introduce a combinatorial notation, which we explain with an example, 2TDM+. In this notation, the first integer (2 in the example) represents the numbering of the state from 1 to 75 (S1 Table). The following characters represent states of the first (red one in Fig 1D) and the second (blue one in Fig 1D) motor domains; in this example the red motor domain is in the T state, while the blue one is in the DM state. The last symbol, either +, 0, or -, represents the relative position of the two motor domains along MT.

### Effect of external forces

Our model incorporates the effect of external force. We suppose the external force is applied to the tip of the linker through the attached bead or cargo in the direction parallel to MT. In this work, we limit ourselves to the case where the external force is applied to the plus end (backward direction to the functional movements) of MT. When one motor domains is ahead of another domain, i.e. the state with + or -, it is natural to assume that the external force is applied only to the motor domain that locates in the opposite side to the direction of the external force. Thus, the external force *F* is applied to the leading motor domain located closer to the minus end. When the two motor domains take the same position along the MT, we assume each motor domain is subjected to a half of the external force *F* / 2.

Since the power-stroke changes the bead position, transition rates for the power-stroke and its reverse process must be affected by the external force. We can easily incorporate this effect into the energy difference between pre- and post-power-stroke states as,

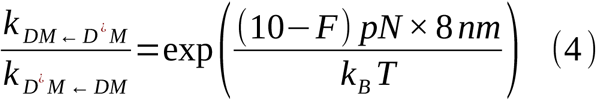

Of note, the external force *F* takes the positive value when dynein is pulled to the plus end direction of MT. Based on this equation, we rescaled the rate of the reverse of the power-stroke process, making it more probable to occur as the force is applied, whereas the rate of the power-stroke is unchanged.

Similarly, the diffusive motions are affected by the external force. It is natural to assume the ratios between *λ*_*for*_ and *λ*_*back*_ depends on the external force as

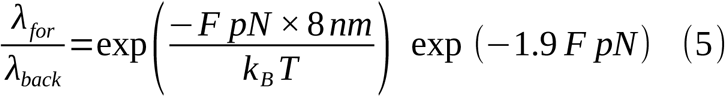

We have multiple choices of *λ*_*for*_ and *λ*_*back*_ that satisfy this equation. Here, we put the force dependence in *λ*_*back*_ making the backward diffusion more probable to occur as the force is applied, while the rate of the forward step is unchanged. Besides, previous experiments uncovered that the dissociation rates of the motor domain from the MT increases with the external force [41,42]. Since this could be an essential feature of the dynein motor domain, we include this effect into our model. Specifically, all the transitions from MT-bound states to unbound states, such as D*M→D and DM→D, that occur in the leading motor domain are accelerated by a force-dependent factor. Note that this corresponds to the increase in the free energy of the leading domain MT-bound state by the logarithm of the force-dependent factor. We introduce a multiplying factor *f* _*ext*_ (*F*) into the dissociation rates of the motor domain under the force *F*. An example is illustrated in Fig 1E. We set *f* _*ext*_ (*F*)=exp (*B · F*) because the associated energy changes should be proportional to the external force. Here, B is a fitting parameter that we estimate below. When the motor domain that is subjected to a half of the external force dissociates from MT, the factor becomes 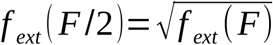 (see an example in Fig 1E from the third to the fourth states).

### Internal force between two motor domains

Since the two motor domains are connected, each motor domain should receive internal force from the other one when the both motor domains are bound on MT. We treat the internal force being independent of the external force. Previous experimental study shows the asymmetric dissociation rates of MTBD from MT for the case of internal force[41,42]. Therefore, the lagging motor domain should dissociate with larger rates than that of the leading (minus-end side) motor domain due to the opposite directions of internal force for the leading and lagging motor domains when both motor domains are bound to MT with 8-nm gap,. We incorporate this effect by introducing a multiplying factor *f* _∫¿¿_ into the dissociation rate of the lagging motor domain when the leading domain is also bound on the MT (Fig 1F).

To satisfy the condition that the free energy change along any cyclic path must be zero, we put the same factor *f* _∫¿¿_, either in the numerator or in the denominator, of many surrounding transition rates, as in S1 Fig. We begin with the acceleration of the transitions by *f* _∫¿¿_; 4D*MDM+ → 24D*MD+ and 1DMDM+ → 21DMD+ (see Fig 1F), in which the lagging motor domains are peeled out due to the inter-motor domain force. These changes are accommodated by increasing the free energies of 4D*MDM+ and 1DMDM+ by ln ¿, which can be viewed as the internal tension. These free energy changes in the two states result in including the same multiplying factor, either in the numerator or in the denominator of all the rate constants connected with these two states (Fig 1F for an example and S1 Fig for the complete picture). For example, in S1 Fig, 4D*MDM+ can transit to 5DDM+ and 9D*MT+ so that we introduce *f* _∫¿¿_into the incoming rates from these two states.

### Setting up the rate constants

Now, we discuss the values of rate constants. As of now, due to the lack of experimental measurements, we cannot decide many parameters without uncertainty. Yet, we would like to choose a set of parameters that satisfy the physics law, equations (1)-(4), and that are consistent with all the available data as much as possible.

First, from single-molecule experiments for the monomeric dynein, we can set

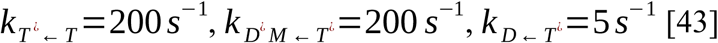

Second, the transition DM→T contains the ADP release 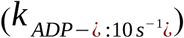, ATP binding 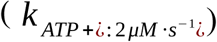, and the dissociation from MT (*k*_*off*_: 500 *s*^-1^), as a roughly sequential process [43]. So we can express *k*_*T* ← *DM*_ as

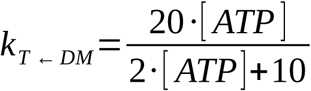

In the derivation process, we assumed that the dissociation from MT is much faster than the other processes. Thus, *k*_*T* ← *D*_ becomes the same as *k*_*T* ← *DM.*_

Third, since direct experimental data are not available at the moment for other values, we infer physically feasible values guided by the detailed balance conditions and other restraints. We assume the power-stroke motion is fast enough and set it as 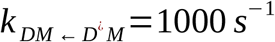. Of note, as far as this is large enough, the result does not depend on this precise value at all. Using the physical constraint, eq.(4), we also can obtain

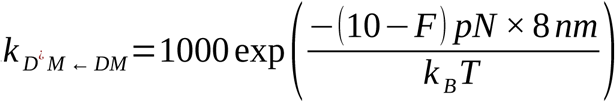

Next, we suppose that the high affinity MTBD does not dissociate from MT without the applied external or internal force. We thus choose *k* _*D*← *DM*_ as a sufficiently small value; specifically, it must be much smaller than *k*_*T* ← DM_ 10 *s*^-1^ at its saturated value. Since both *k* _*D*← *DM*_ and 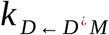 represent dissociation of the high affinity MTBD from MT, we set 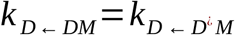. The D state must be inherently unstable. Thus, the lifetime of the D state is short and it changes quickly to the DM state. It is natural to assume *k*_*DM ← D*_ >*k*_*T ← DM*_10 *s*^-1^. From the detailed balance condition, we get

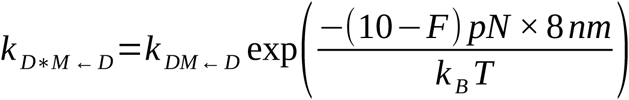

Others rate constant values are determined to satisfy the detailed balance. Finally, *k*_*T* ←*T*\*¿¿_ should be smaller than the reverse process, *k*_*T* *←*T*_.

We list all the kinetic parameters used in the current simulations in Table 1.

**Table 1.**
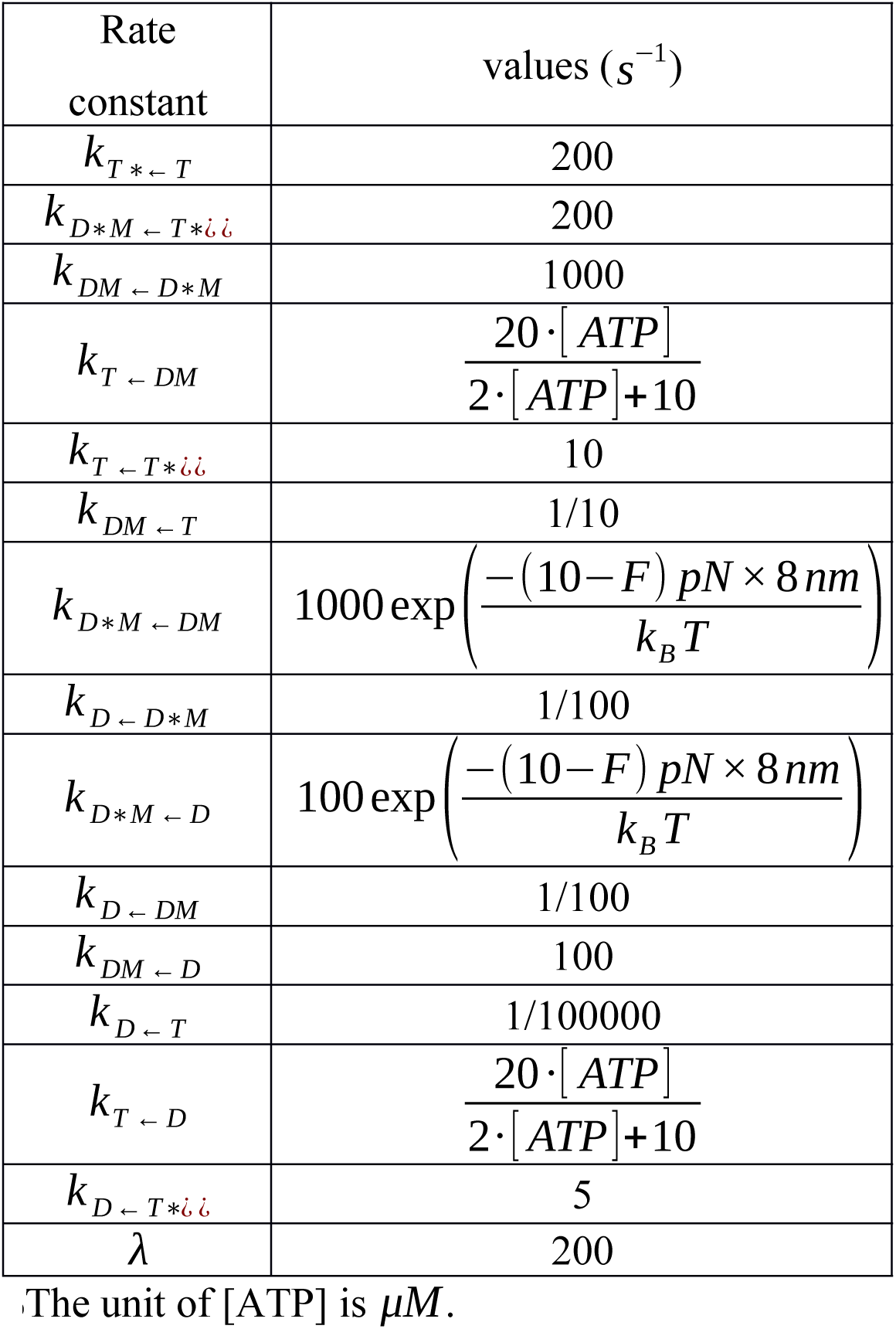
Rate constant values for motor domain.

Next, we determine the external force dependent factors. The detachment force of dynein, i.e., a critical force beyond which dynein detaches from MT promptly, to the plus end of MT with the high affinity MTBD is estimated to be approximately 2 pN (Shima et al, manuscript in preparation). This suggests that, with the external force of 2 pN, the DM state should change to the D state, rather than the other states, i.e., the T state. Thus, we require

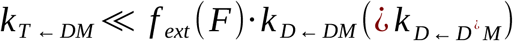. We note that *k*_*T* ← *DM*_ =10 *s*^*-1*^ at the saturated ATP concentration. Since we set the dissociation rates with no external force to be 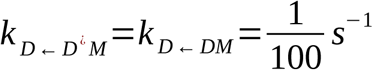, previously, we decided to choose *f* _*ext*_ (2 *pN*)=10000. To this end, we obtain *f*_*ext*_(*F*)=100^(*F pN*)^ exp (4.61 *F pN*). Note that the estimate of the detachment force contains some uncertainty; if the detachment force were equal to 3 pN, we would have *f* _*ext*_ (*F*) exp (3.07 *F pN*)

While there is no experimental estimate for the value of *f* _∫¿¿_, merely for the consistency with the effect of the external force in the previous section, we set *f* _∫ ¿=10000 ¿_; we suppose the dissociate rate of the relevant motor domain from MT may be equal. We also tested how this value would alter the results, finding that most of the qualitative results are not changed, except the probabilities to choose individual paths (see Discussions).

### Monte Carlo simulations

As the basic dynamic equations for the kinetic model of 5 ×5 ×3=75 states, we take the master equation;

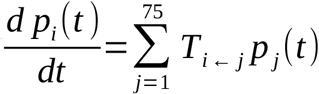 where *p* _*j*_(*t*) is the time-dependent probability to be in the j-th state and the transition matrix *T* _*i*← *j*_ represents corresponding transition rates for the non-diagonal elements and cumulative outgoing rates for the diagonal elements, respectively. Of the 7 5 ×75 matrix elements, we have already described all the non-zero elements, while the other elements are zero. In this work, we solved this master equation via Gillespie Monte Carlo (MC) simulations, a random number-based sampling of the solution. When the system resides in j-th state at a time t, the Gillespie algorithm first chooses the state to go; among those connected from the current state j, a state i is chosen by the probability proportional to the corresponding transition rate *T* _*i*← *j*_. Next, using the summation of all the rates connected from the state j, 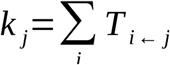, the transition time Δ *t* is drawn randomly from the waiting time distribution in Poisson process, *P*(Δ *t*)=*k* _*j*_ exp (-*k*_*j*_ Δ *t*). In all the MC trajectories, we begin with the initial state, 1DMDM+, the red motor domain in the DM state is ahead of the blue motor domain, which is also in the DM state. Each MC simulation is terminated at the moment when both motor domains are disassociated from MT (i.e., T, T*, or D state) or when the time exceeds 10s. For each setup, we repeated MC simulation 10000 times with different random number sequences.

We also note that the current master equation can be solved by the standard linear algebra, as well. We tried to solve the linear algebraic equation by an algebraic manipulation software. However, we did not succeed to obtain the closed formula for this large matrix. Still, one can solve the linear equation numerically, obtaining the exact numerical values for velocity and the run length. However, the MC simulation is more straightforward to analyze pathways, and thus we took the MC approach in this study.

### Analysis of the run length from trajectory

When we determine the run-length from a trajectory, to avoid possible artefacts in the initial configuration, we used a scheme used in experiments[38]. First, we plotted the cumulative probability distribution *c* (*x*) (red crosses in Fig 2B) as a function of the run length *x*, and confirmed that 1-*c* (*x*) exhibits exponential decay for *x* >1step. This is probably due to the Poisson process of the dynein detachment. Given the exponential behavior, we fitted *c* (*x*) with the form,

**Fig 2.**
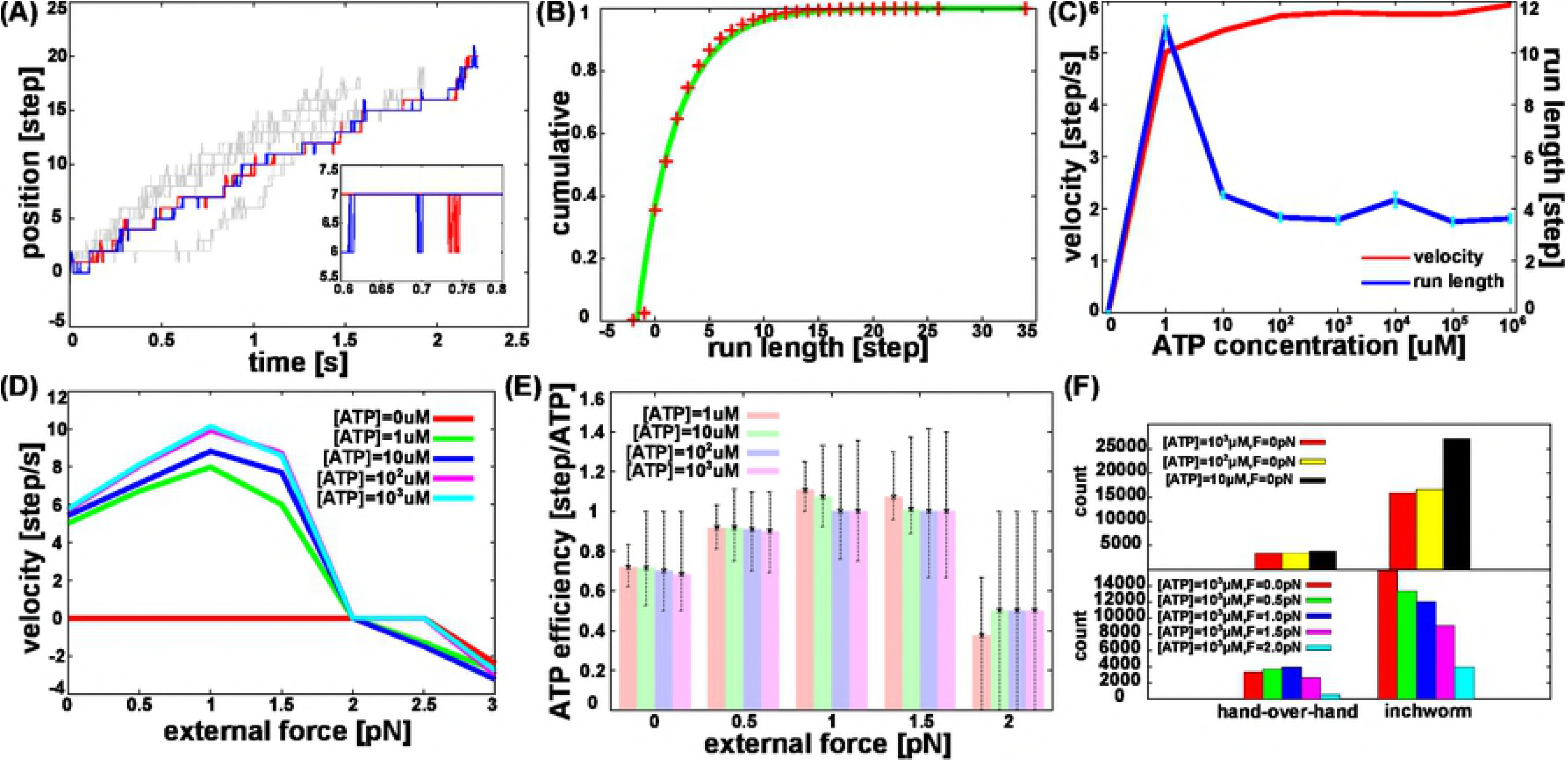
Bipedal motions of dimeric dynein via Monte Carlo simulations. (A) Five representative trajectories at [ATP]= 1mM with no external force. The red and blue curves, respectively, represent positions of red and blue motor domains along MT in one trajectory. Grey curves represent movements of two motor domains in four other trajectories. The inset is a close up between 0.6s and 0.8s. (B) The cumulative probability distribution (red crosses) calculated from the final arrival distance for 10000 MC trajectories under the same condition as (A) and its fitted curve (green). (C) Median velocity and mean run length as a function of [ATP] with no external force. Error bars from each point of *x*_*mean*_ are asymptotic standard errors. (D) Mean velocity as a function of the external force for a few different [ATP]. (E) Median number of steps per consumed ATP (ATP efficiency) for a few different [ATP] and external forces. The error bar represents quartile range. (F) Histograms of the hand-over-hand and inchworm motions.

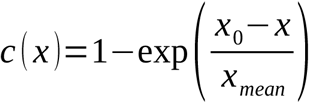

to obtain *x*_0_ and the mean run length *x*_*mean*_ by the nonlinear least square estimation with the Marquardt-Levenberg algorithm. Here, *x*_0_ is used to accommodate irregular behavior of the very initial stage.

## Results

### Kinetic model reproduces many data on the wild-type dynein motility

First, we examine the basic motility of our kinetic model comparing the simulation results with experimental data. We performed MC simulations of wild type dimeric dynein movement with [ATP] = 1 mM and no external force. We repeated simulations 10,000 times, from which five representative trajectories are shown in Fig 2A (see S2 Fig for the entire trajectories). For this trajectory, we see that dynein stochastically proceeded to the minus-end direction of MT for ∼20 steps before detachment from MT at ∼2.2s. Interestingly, we find dynein sometime moved quite rapidly, but also exhibited occasional pauses. During the pauses, we find some rapid stamping of one motor domain (while one domain pauses in a position, the other domain moves back and forth many times, as in the inset of Fig 2A). The other four trajectories (drawn in grey curves in the figure) proceeded to the same direction and roughly with the same velocity.

From the 10,000 trajectories, we estimated the mean run length; for example, for the case of [ATP] = 1 mM, it was estimated as 3.6 *±* 0.15step (Fig 2B). We plot the mean run length as a function of the ATP concentration [ATP] (Fig 2C). The mean run length shows a peak at around [ATP] = 1μM and levels off at higher [ATP]. The ATP-bound dynein, the T or T* states, has weak affinity to MT and thus the saturated [ATP] diminish the run length. The characteristic event that the run-length becomes longer with lower [ATP] was also reported in 22S dynein of Tetrahymena cilia and cytoplasmic dynein of mammalian [44,45].

We then plot the velocity as a function of the ATP concentration [ATP] (Fig 2C). For the velocity, we defined it as the ratio of the distance of movement to the time duration between long-time pauses, from which we obtained the median *v*_*med*_ of the velocity distribution as the representative value. We note there can be several other ways to define the velocity, which we discuss in the supporting information (S3 Fig). As [ATP] increases, the velocity increases at the low [ATP] and then saturates at higher [ATP], which is qualitatively consistent with experiments. In the saturated [ATP], each motor domain is bound by ATP for most of time and the ATP binding does not determine the velocity, as usual. The maximum velocity in the model dynein was about 6 step/s ∼ 50 nm/s.

Experimentally, the velocity of dynein movement has been measured and known to depend on the species/constructs. The velocities for the full length and the GST dimer of *Dictyostelium* dynein are ∼200 nm/s [46], and ∼500 nm/s [47], respectively. For mammalian dynein, the DDB complex proceeds with 500∼800 nm/s and the GST dimer velocity is ∼500 nm/s [48]. For yeast, the full length dynein moves at the velocity ∼80 nm/s [8] and the GST dimer proceeds with ∼100 nm/s [49]. Our model dynein moved with the velocity similar to that of yeast dynein Reasons that the model shows lower velocity than *Dictyostelium* dynein may be attributed to the choice of the low rate constant for *k*_*T* ← *DM*_ which actually could dramatically change depending on the dimeric state [43].

Next, we calculated dynein motions under the external force to the backward direction, i.e., towards the plus-end of MT. The estimated velocity *v*_*med*_ is plotted against the strength of the external force for a few different [ATP] in Fig 2D. With non-zero [ATP], the velocity is positive at low external force and becomes negative at sufficiently large force, as expected. Notably, the mean velocity crosses zero at the same external force (∼2 pN), regardless of [ATP]. This is in harmony with a recent experimental result in dynein motility assays (Shima et al, manuscript in preparation). Another non-trivial behavior is a slight increase in the velocity with a weak external force of 0 - 1 pN, which we will discuss later.

We also estimated the efficiency of our model dynein. Specifically, we plotted in Fig 2E the median forward step numbers per one ATP hydrolysis, termed ATP efficiency for brevity, for several combinations of [ATP] and the external force. Focusing on the ATP efficiency in the absence of external force, although the run length and the velocity differ by far between [ATP]= 1 *μM* and [ATP]= 10^3^ *μM,* the ATP efficiency changes less than 5%. Namely, the average move per one ATP cycle does not depend on the ATP concentration much, whereas the waiting time for ATP binding depends on [ATP] and thus affects the velocity at low-to-medium range of [ATP]. As [ATP] increases, the ATP efficiency decreases slightly from 0.71 at 1 *μM*, to 0.68 at 10^3^ *μM.* This is because the forward move by the power-stroke is partly canceled by the influence of diffusion in the weakly coupled dissociation state. When we applied the external force to the backward direction, we found the increase in the ATP efficiency for the range [ATP] = 1-10^3^ *μM* and up to the force of 1pN (the efficiency reaches to ∼1). Above 1.5 pN, the efficiency decreased. This is related to the increase in the velocity with a weak external force mentioned above, and we will come back to this feature in the pathway analysis later.

Notably, with large external forces (1.5 pN and above for [ATP] = 1 μM, and 2.0 pN for the other [ATP]), we found a much larger variance in the ATP efficiency than the cases of weaker force. With large opposing force, dynein occasionally goes backwards largely and then detaches from MT, which makes sampling of broad data difficult.

### Mutants that impair one motor domain activity

It is an interesting feature of dimeric dynein that it can proceeds to the same direction even when one of motor domains lacks the ATP binding or hydrolyzing ability[11,50]. Here, to test our model, we performed MC simulations for the two cases that mimic the two types of impaired heterodimeric mutants. Fig 3A shows representative trajectories for the case where one motor domain does not bind ATP. Clearly, with the ATP-binding deficient mutant in one motor domain, it still moved to the forward direction with qualitatively similar processivity. Not surprisingly, during the processive movement, the intact motor domain was ahead of the mutant motor domain in most steps. We also see this mutant showed slightly reduced velocity and slightly increased run length, compared to the wildtype (grey in the figure) (Fig 3C and 3D). This is because the mutation leads to increase the population in the high-affinity state to MT, which slowed the movement and increased the processivity. Both of these effects have been shown in previous experiments, suggesting our model calculation qualitatively agreed with the experiments[50].

**Fig 3.**
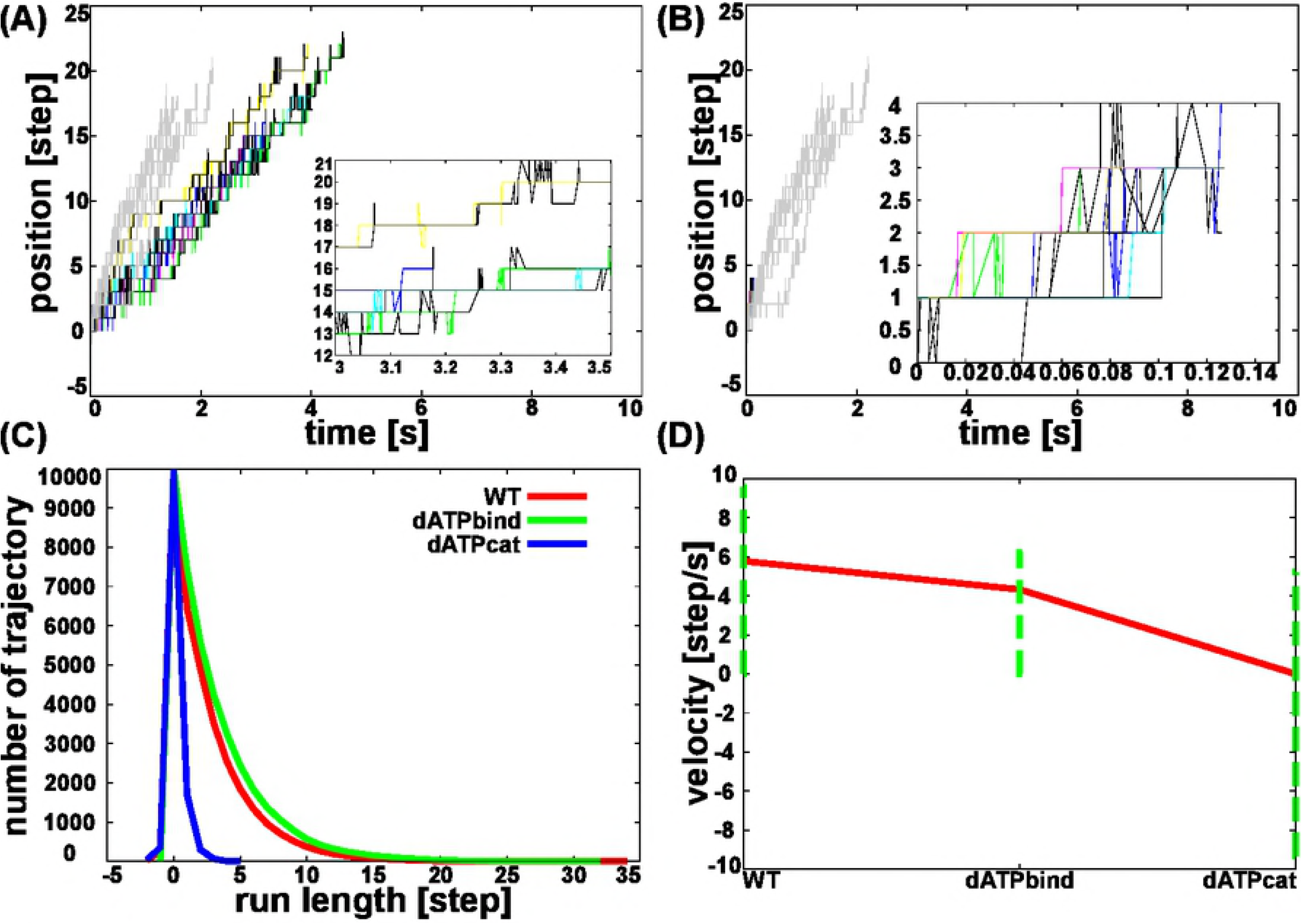
Motility of two mutants of dynein. (A) Representative trajectories at [ATP] = 1mM with no external force for a mutant of which right (blue) motor domain does not bind ATP. The positions of the deficient motor domain are drawn in black. Grey trajectories are for wild-type. The inset is a close up between 3s and 3.5s. (B) Representative trajectories at [ATP] = 1 mM with no external force for a mutant of which right (blue) motor domain does no hydrolyze ATP. The positions of right motor domain are drawn in black. Grey trajectories are for wild-type. The inset is a close-up view from 0s to 0.14s. (C) Histogram of the run length. Red, green, and blue curves represent the wild-type, the case where one motor domain does not bind ATP, and the case where one motor domain does not hydrolyze ATP, respectively. (D) Mean velocities and their standard deviations for the three cases as in (C).

Fig 3B plots trajectories for the case where one of motor domain does not hydrolyze ATP. The ATP-hydrolysis deficient mutant moved forward, but markedly reduced the processivity and run length because the mutated motor domain stay in T or T* states with low affinity to MT. We also see this mutant showed slightly reduced velocity, compared to the wild-type case (grey in the figure) (Fig 3C and 3D). Indeed, previous experiments showed decreases in the run-length and the velocity for this mutant [50]. Thus, our model calculation qualitatively agrees with the experiment.

We note, however, that experiments showed the ATP-hydrolysis deficient mutant moves with larger velocity than the ATP-binding deficient mutant, which differs from our model calculations. We consider that our model dynein of the ATP-hydrolysis deficient mutant detaches from MT quickly so that it is difficult to complete ATP cycles in many trajectories, which makes the velocity estimate difficult.

### The bipedal mechanism: Pathway analysis

So far, we described overall behavior of motility in our kinetic model, showing that the model can reproduce many of previous experimental observations. Now, we analyze the underlying mechanisms emerged from our model. For this purpose, we focus on the case of [ATP] = 1 mM with no external force, unless otherwise denoted.

Fig 4 shows the whole network of the states-to-state transition dynamics, which, albeit of high complexity, contains full of mechanistic information. Hereafter, we decipher this complex network. Of the 75 states, the network contains only those that appeared in our 10000 trajectories.

**Fig 4:**
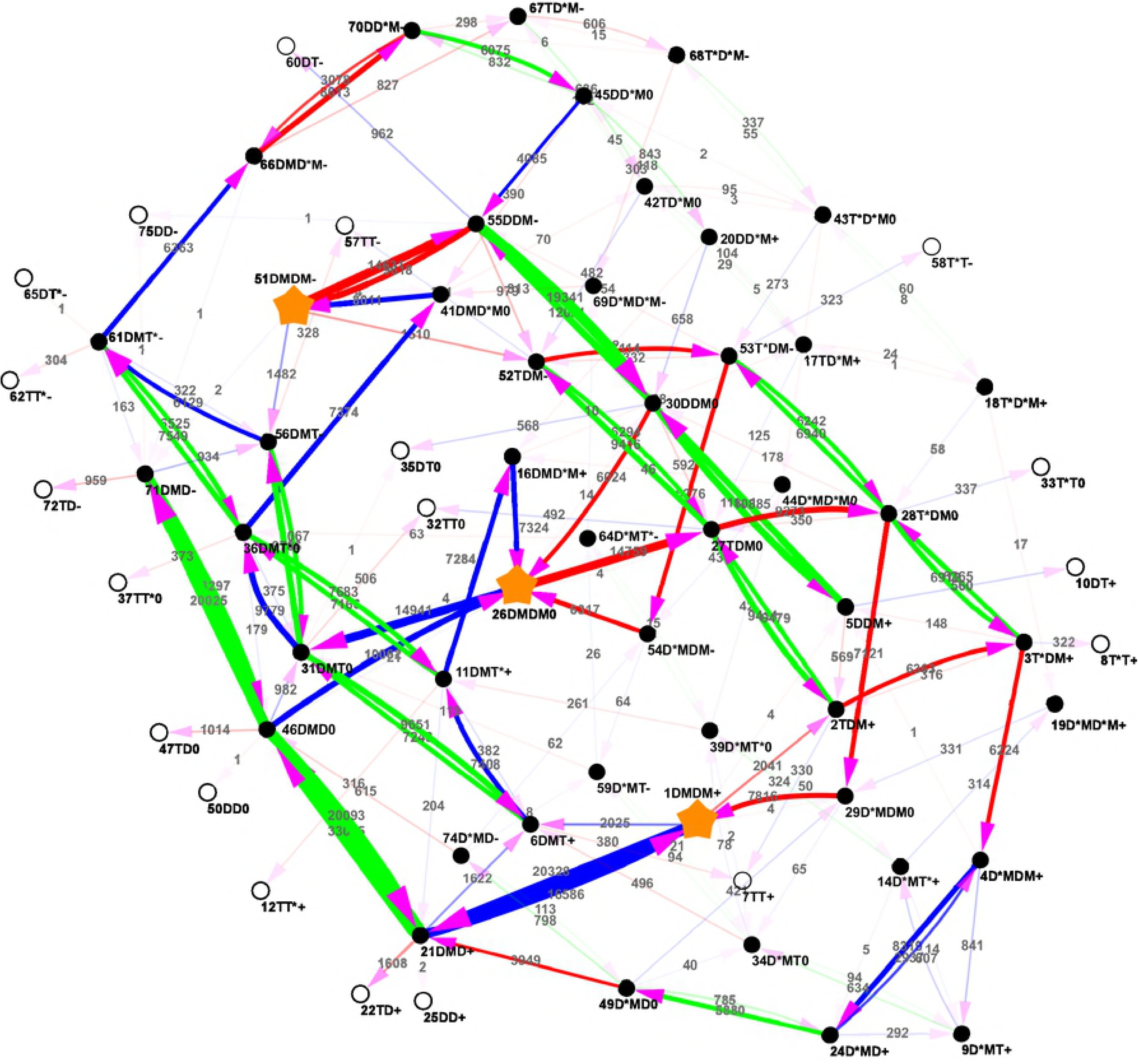
The whole network of pathways. Black filled and open circles represent the dimeric dynein states bound on and detached from MT (the dead-end), respectively. Red (blue) arrows indicate the transitions where the state in the red (blue) monomer changes, whereas the green arrows mean the diffusive motions along MT. Integers written on the arrows represent the number of transition times observed in simulations, with which the thickness of the arrow correlates. For the meaning of the label of each state, see the text. 1DMDM+, 26DMDM 0, and 51DMDM - are marked with stars, to emphasize their high populations.

Of the 75 states, the most populated states were the 26DMDM0 state (see S2 Table for the list of high population states); both monomers are in the DM states located at the same position along MT (implicitly, bound on different proto-filaments) (a star mark located near the center of the figure).

Starting from this ground state 26DMDM0 and following probable transitions (thick arrows, with the counts larger than 5000), we find a pair of probable cyclic pathways. The one that makes a clockwise rotation in the upper side of the figure is

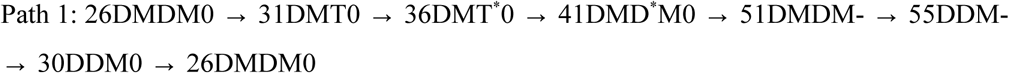

On this pathway, from the 26DMDM0 state to the 51DMDM-state via three transient states, the blue motor domain completed one ATP hydrolysis cycle while the red one remained in the DM state, by which the blue motor domain proceeded by one step forward via recovery-stroke and the subsequent power-stroke of the linker (Remember that, to distinguish two identical motor domains, we call the first and the second domains as the red and the blue domains, respectively). The 51DMDM-state is a long-lived intermediate state (star-marked in the figure). Subsequently, the lagging red motor domain detached from MT reaching to 55DDM-, which is followed by the diffusive motion of the red motor domain 30DDM0. Finally, the red domain rebound to the MT returning into the starting state 26DMDM0. When the dimeric dynein repeats this cycle more than once, this process is normally called the inchworm motion. Notably, during this cycle, the ATP hydrolysis reaction occurred only in the leading (blue) motor domain, whereas the lagging (red) motor domain is simply dragged by the leading one. This is in harmony with the observation that the mutants that impair motor activity still move in one direction. The other prominent cycle (a clockwise cycle in the bottom side of the figure), which is related to the first one by symmetry, is,

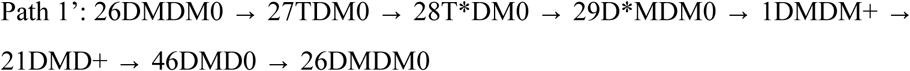

in which the red motor domain hydrolyzes one ATP to drive the system forward. Clearly, the mechanism is identical to the first cycle.

Thus, while the pathway analysis exhibit extremely diverse routes, we find a dominant cyclic pathway which corresponds to the inchworm motion.

### Distinct bipedal motions

While we found one prominent cyclic pathway, Path 1, in the previous section, due to the complexity of the whole network in Fig 4, we need a more systematic analysis to reveal various pathways. Given that each motor domain has the highest population in the DM state (S2 Table), we systematically seek pathways that connect the dimeric dynein states where both motor domains take the DM states. Among several possible combinations, we found the two cases are dominant; 1) the one starting from and ending to the 26DMDM0 state and 2) the other starting from 1DMDM+ and ending at 51DMDM-without passing through the 26DMDM0 state. We note that, by symmetry, we also observed the case from 51DMDM- to 1DMDM+, of which the mechanisms are identical to the second case.

We depict the schematic pictures of the major cyclic paths starting from and ending at the 26DMDM0 state in Fig 5A. Clearly, all these correspond to the inchworm motions when the cycles are repeated more than once. In the 10000 trajectories, we identified 15896 steps of inch-worm motions. The figure contains, in the second row, the Path 1 already described above based on the visual inspection.

**Fig 5:**
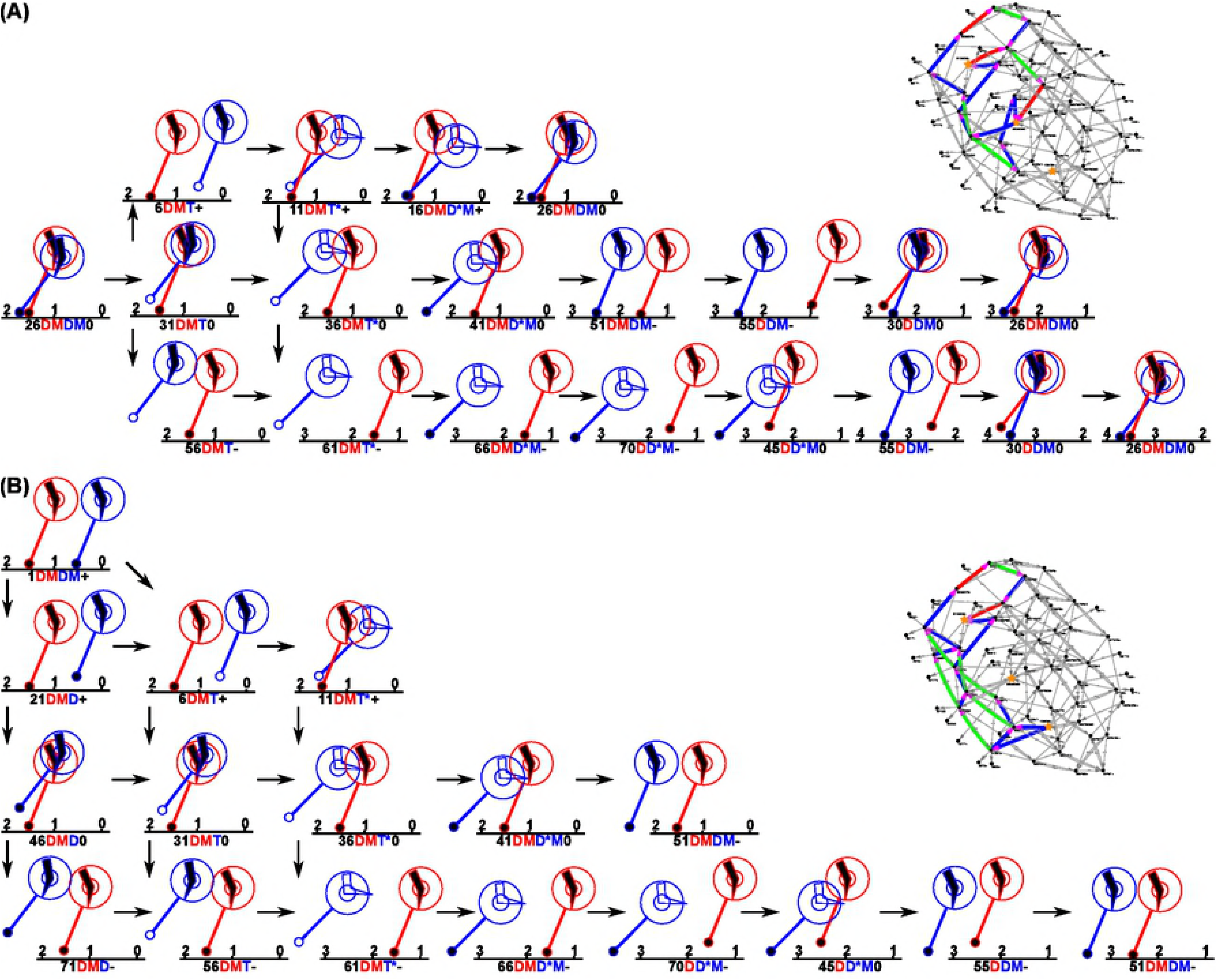
The two distinct bipedal working pathways. (A) The inchworm-like pathway. (B) The hand-over-hand pathway. Schematic diagrams and the whole network are drawn on the left and the right, respectively.

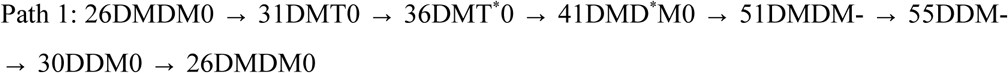

As described above, the leading motor domain made one ATP hydrolysis cycle driving this domain forward by one step, whereas the lagging motor domain moved via diffusive motions dragged by the leading domain. In addition, we find a branch path depicted in the top row of the figure, in which the detached leading motor domain diffused backward, which is followed by the recovery-stroke and subsequently the power-stroke. Overall, the molecule returned to its original state and position. Moreover, we find another branch path depicted in the bottom line of the figure, where the detached blue motor domain diffused forward. The following steps include recovery-stroke and then the power-stroke in the blue motor domain. Together, the dynein moved 2 steps forward, one by diffusion and the other by the power stroke. Thus, while a pair of linker recovery-stroke and the subsequent power-stroke in the leading domain contributes to the one forward step, additional diffusive motions, if any, can modulate the motions.

Next, we consider the paths starting from 1DMDM+ and ending at 51DMDM-without passing through the 26DMDM0 state, of which a schematic picture is drawn in Fig 5B. Albeit high diversity, these paths all correspond to the hand-over-hand motions. In the 10000 trajectories, we identified 3328 steps of the hand-over-hand motions, which is about one fifth of the inch-worm motions (Fig 2F). This route contains many branches, but each path includes one or more relatively slow transition processes, making this entire route not as frequently used as the Path 1.

Among the many branches during the hand-over-hand motion, the most frequently used path was as follows,

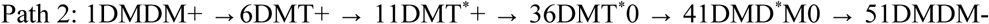

During the process, fueled by the ATP hydrolysis reaction cycle in the blue motor domain, the blue domain moved from the rear side of the red motor domain to the forward side by 2 steps, and thus 16 nm. We note that one of the 2 steps is via diffusive movement. More in details, starting from the 1DMDM+ state, the internal tension from the forward red motor domain induces the ADP dissociation/ ATP binding in the rear blue domain, leading to the detachment from the MT (6DMT+) and subsequently recovery stroke (36DMT*+). Then, the lagging blue domain diffuses forward (36DMT*0), which is followed by the ATP hydrolysis (41DMD*M0). Finally, the power-stroke in the blue domain moved the blue domain one more forward step. We note that in this process the one (blue) motor domain moved by two steps, one via diffusion and the other by the power-stroke. The other (red) motor domain remained in the DM state through-out. Interestingly, the hand-over-hand motion in the model dynein is qualitatively different from that of kinesin-1 (or conventional kinesin), the neck-linker docking (classically termed the power-stroke) occurs in the MT-bound leading head. Whereas, our model dynein uses the power-stroke of the lagging motor domain to move itself forward. When the dimeric dynein continues on this bipedal motion, the next step is that the red motor domain moves forward by two steps using ATP hydrolysis reaction. Therefore, mutants that impair one motor domain activity cannot move by this mechanism. There exist similar paths closely related to the Path 2, such as

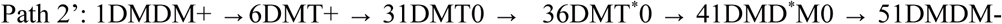

which differs from the Path 2 only in the third state 31DMT0. These are essentially the same. Occasionally, we observed that both the blue and red motor domains made diffusional move-ment during one ATP cycle in the blue domain. The path can be described as

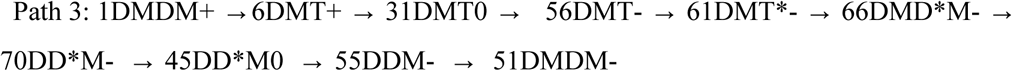

This whole path contains three diffusive movements, in addition to one ATP hydrolysis cycle in the blue motor domain.

In summary, starting from the states where both motor domains are in the most stable DM states, we found two prominent cyclic pathways, Paths 1 and 2, which correspond to the inch-worm and the hand-over-hand motions, respectively. By counting the respective cycles in the trajectories, we found that the inchworm motions are more probable than the hand-over-hand motions for all the [ATP] and the external force conditions tested here (Fig 2F). The predomi-nance of the inchworm steps in the model dynein is in harmony with the experiment data [10].

### Fast-track, slow-track, and back-step

So far we described the inchworm and the hand-over-hand motions in the model dynein stepping, but the whole network includes many more sub-dominant pathways. To address effects of dynein stepping pathway in its velocity, we classified fragments of trajectories by the short-term velocity defined within the fragment. Here, we define the three classes, fast-track, slow-track, and backward move and discuss dominant pathways in each class (we exclude the medium-velocity class because it corresponds to the dominant pathways discussed above). The fast-track is defined as the fragment of trajectories that have their velocities faster than the third quartile value (480 nm/s). Similarly, the slow-track is defined as those with the velocities slower than the first quartile value and that are positive (54 nm/s). The backward move is defined as the negative velocities in the fragment of trajectories.

First, Fig 6A depicts the fast-track pathways and its schematic mechanism. One prominent path starts from the 1DMDM+ state (or equivalently 51DMDM-state) and proceed via the Path 2 (the hand-over-hand manner).

**Fig 6.**
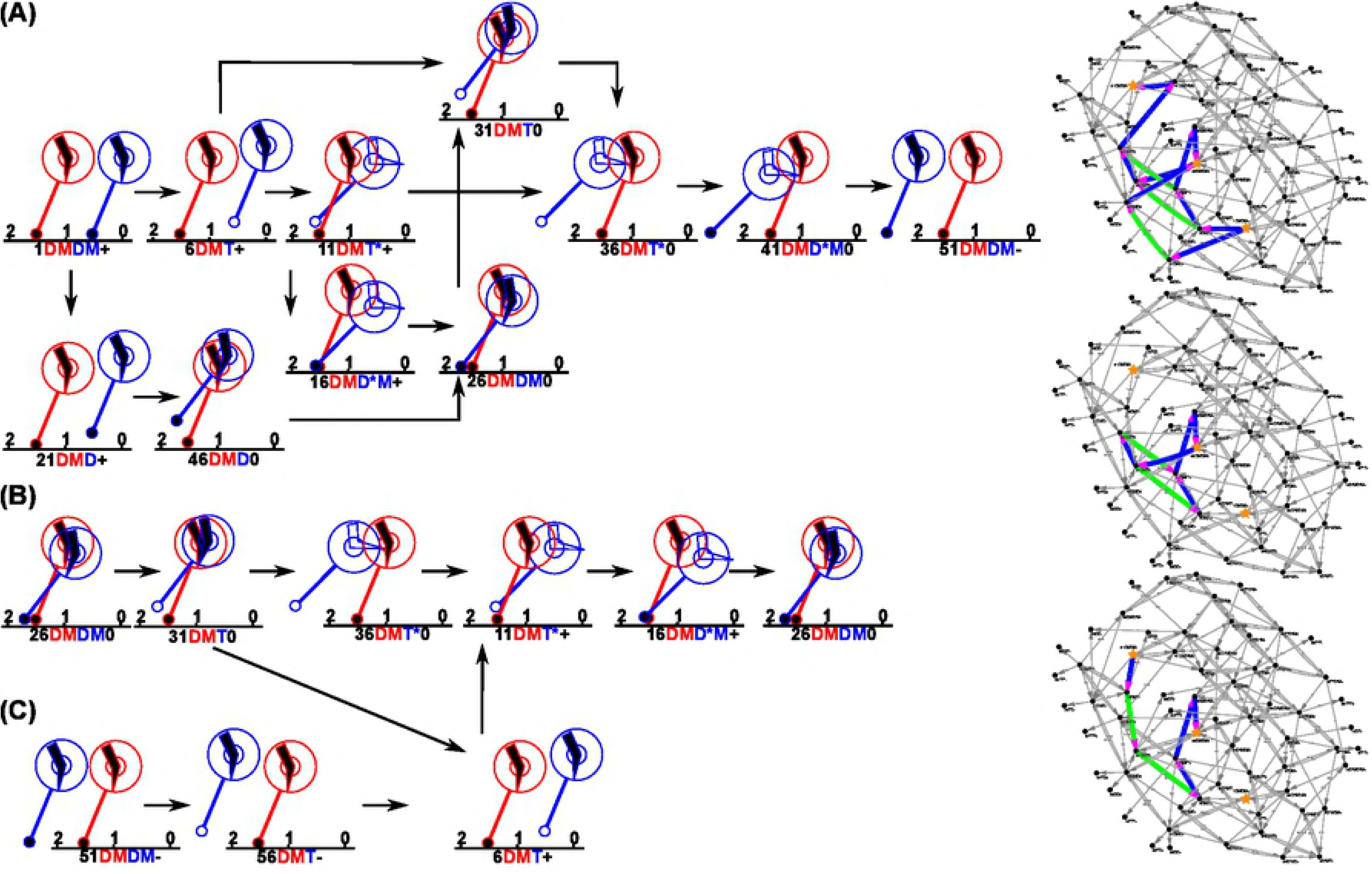
Dominant motions in different velocity regimes. (A) Fast-track motion. (B) Slow-track motion. (C) Backward motion. Schematic diagrams and the whole network are drawn on the left and the right, respectively.

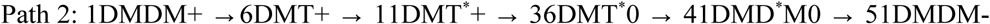

As above, in the fast-track pathways, there exist some small variant paths. For example, when the order of the recovery stroke and the diffusion is exchanged from the above path, we obtain a variant path depicted at the top line in the left side of Fig 6A. Alternatively, starting from the initial 1DMDM+ state, detachment and the diffusion of the rear domain can precede the nucleo-tide exchange reaction, which is drawn at the bottom line of the left side of Fig 6A. In summary, the hand-over-hand motion dominates the fast movement.

Next, we discuss the slow-track, of which schematic pathways are drawn in Fig 6B. The prominent pathway observed is

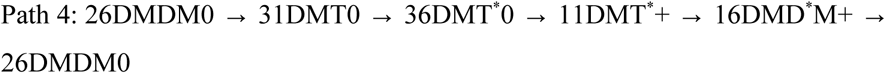

The first three states in this path are identical to that in the Path 1. From the 36DMT^*^0 state, the blue motor domain diffused backward (11DMT^*^+). Then, the blue domain was bound on the MT upon ATP hydrolysis (16DMD^*^M+), which is followed by the power-stroke to return the 26DMDM0 state. This path was illustrated in Fig 2A at around the time 0.70 - 0.74 s where we see long pause with an instantaneous back step of the blue domain. Overall, the molecule did not proceed forward, but came back to its original location. A variant of the Path 4 is also depicted at the bottom line of the left side of Fig. 6B. Pathways observed in the low velocity class always include a backward diffusion.

Finally, we describe the case in which the dimeric dynein moved backward. Fig 6C merged with Fig 6B represent a pathway,

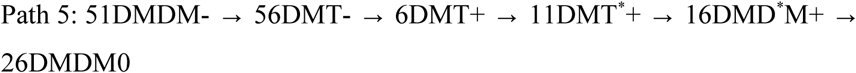

The last three states in this path are identical to those in Path 4. A crucial feature in this pathway is that, in the very first transition, the leading blue domain detached from MT, which is in contrast to the detachment of the lagging domain in the Path 2; the hand-over-hand motion. Once the leading blue domain is detached, it cannot diffuse forward, but can diffuse backward, due to the restraint from the MT-bound red motor domain. In the Path 5, the blue domain diffused backward by two steps, which is followed by the recovery stroke and then the power-stroke motion in the blue domain. Overall, the dimeric dynein moved backward by one step.

## Discussion

### The observed primary inchworm motion uses one ATP per a dimer step

Here, we propose the kinetic model, which can reproduce extremely versatile modes of dynein movement. Our model suggests that the dominant pathway is the inchworm motion, which is about 5 times as many as the hand-over-hand motions. This is consistent with a recent report that classifies the modes of movements of dynein finding that about 80% of steps are of the inchworm type motions [10].

Notably, among some variants, the most prominent inchworm pathway in our model dynein,

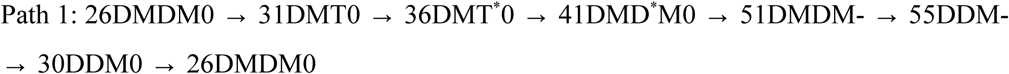

uses only one ATP hydrolysis per a dimer step. Namely, only the leading motor domain moves via ATP-dependent linker power-stroke coupled with a change in the stalk angle, whereas the lagging motor domain is moved via diffusion dragged by the leading motor domain without the ATP hydrolysis cycle. In this sense, the observed model is clearly different from previously suggested inchworm models where two ATP hydrolysis are assumed to occur per a dimer step (e.g., Fig. 4 in [51], Fig. 2 in [2]). It should be noted that the current model do include such a pathway. In Fig. 3, we find the transition from 51DMDM-to 52TDM-, which continues to make the following cycle,

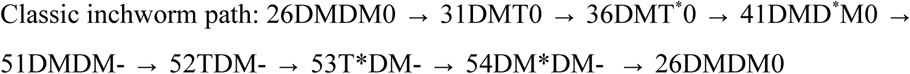

where each motor domain hydrolyzes one ATP, resulting in two ATP consumption per a dimer step. In our model, this pathway is minor since the transition from 51DMDM- to 52TDM-occurs with a relatively low probability. A recent single-molecule assay measured the AAA+ ring angles relative to MT, finding that about 50% of motor domain steps are coupled with small angle changes in the AAA+ ring [52]. The Path 1 is perfectly in harmony with this result because only the leading motor domain takes the ATP hydrolysis cycle which exerts the force and leads to changes in the AAA+ ring angle. On the other hand, the classic inchworm path is not compatible with this single-molecule assay data because it contains twice of ATP hydrolysis cycles and steps and thus it takes 100% coupling between the stepping and the angle change.

Within our model, we assume that the stalk lean towards the MT axis by about 15 degree in the pre-power-stroke state, T* and D*M states, relative to the post-power-stroke state. However, the lifetimes of the pre-power-stroke states are rather short: From the populations in every states (S2 Table), we estimated the probability to have at least one pre-power-stroke state is only 15%, as a whole. Thus, due to short lifetimes of the pre-power-stroke states, some of these angle changes in the stalk might not been observed by the single-molecule assay [53].

It is interesting to discuss an analogy to other molecular motors that are known to use the inchworm motions: Some helicases and ATP-dependent chromatin remodelers are known to use the inchworm motions to proceed along DNA [54–56]. The inchworm motions in these molecules are realized by two subdomains where the ATP hydrolysis is catalyzed at the interface of the two subdomains. In the apo-state, the subdomains are bound on DNA in an open form. Upon ATP binding, subdomains close by sliding one of the two subdomains on DNA by one basepair. After ATP hydrolysis, the subdomains return to the open form by moving the other subdomain on DNA by one base-pair, which results in the one base-pair inchworm motion. Notably, one ATP hydrolysis cycle is sufficient to achive one inchworm motion in these cases. Thus, this usage of ATP is similar to the inchworm motions found in the current simulations for the model dynein.

### The observed hand-over-hand motion agrees with the previous models

We observed the hand-over-hand motions as a sub-dominant pathway. Even though there exist many branch paths, a prominent one is

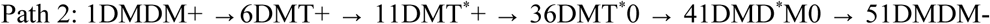

which is actually identical to the hypothetical model suggested previously [11,53]. In our kinetic model, however, this mode is sub-dominant.

It is interesting to note that the hand-over-hand motion in the dynein is distinct from that in kinesin. In kinesin, chemical events in both heads are more tightly coordinated. Starting from the two-head bound states, the ATP hydrolysis in the lagging head reduces its binding affinity to MT and results in the dissociation from the MT. The ATP-dependent neck-linker docking (classically called neck-linker power-stroke) occurs in the originally leading head. On the other hand, the hand-over-hand motion found in our model dynein contains the full of ATP-hydrolysis cycle in the originally lagging motor domain, while the originally leading domain stays in the ADP-bound state. The ATP-dependent power-stroke occurs in the originally lagging motor domain.

### Bipedal motion under external force

It has been suggested that a load exerted by bound cargos speeds up dynein movements by acti-vating some conformational changed. However, our model propose a possibility that this increase in the velocity under load is an intrinsic feature of the dynein motor domain. As in Fig 2D, the velocity slightly increased with a weak external force; e.g., at [ATP] = 1 mM, the velocity with 0.5 pN external force was larger than that without external force. This slight increase in the velocity under a weak opposing force was also reported in previous experimental studies [40,57]. The load-dependent cancellation of dynein auto-inhibition has been proposed for the cause of the phenomenon, however, the molecular basis for such cancellation was still obscure. Our model analysis may partly explain the reason for acceleration of dynein motility under a small backward load.

In order to understand the underlying mechanism of this increase, we sought the difference in the paths with and without 0.5 pN of external force. We extracted fragments of paths that are significantly more frequent with the 0.5 pN force compared to the case of no force (Path 6, depicted in Fig 7)

**Fig 7:**
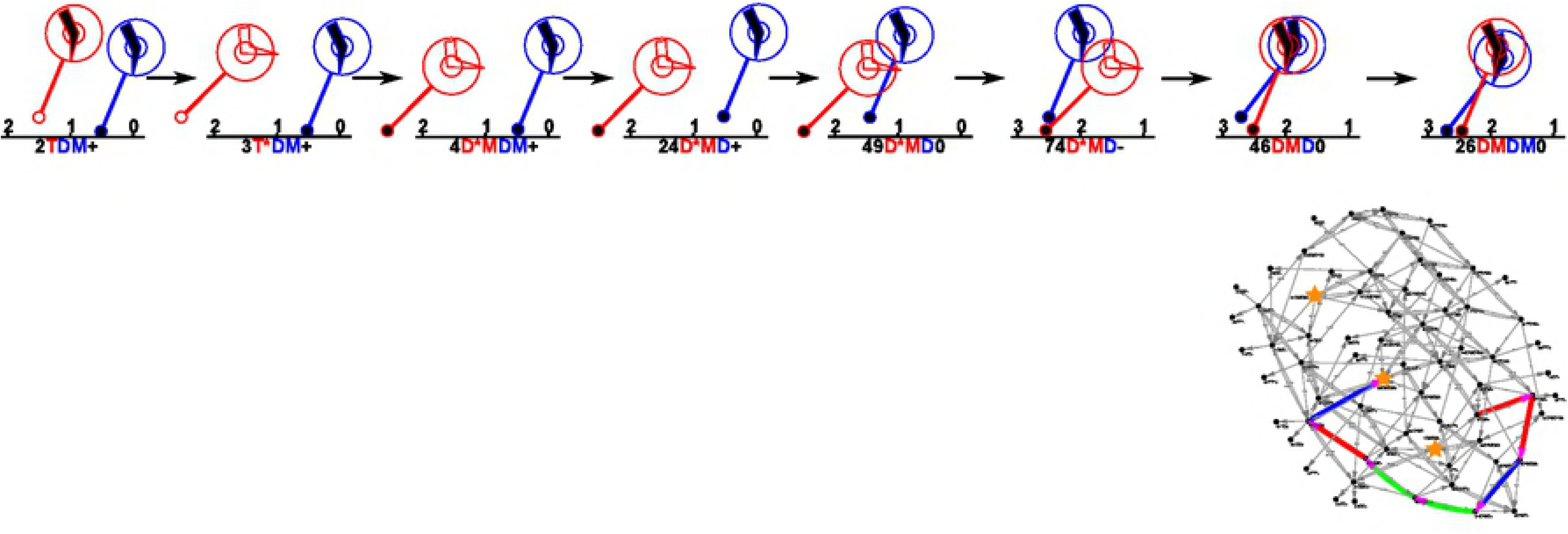
Paths observed in the case of 0.5pN external force more frequently than the case of no external force. Schematic diagrams and the whole network are drawn on the left and the right, respectively.

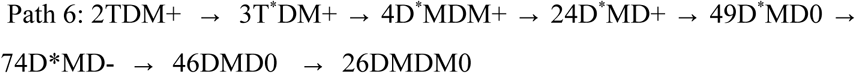

In the first half of the path 6, the leading red motor domain takes the ATP hydrolysis reaction, which inevitably peels out the lagging motor domain from MT due to the internal force. We then ask why this path was enhanced by the 0.5pN force. First, total incoming flows to 2TDM+ were equally probable with and without the external force. Second, the transition from 2TDM+ to 3T*DM+ was notably enhanced by the external force. Without the external force, the 2TDM+ state tends to make a transition to 2TDM0 via backward diffusion of the red domain. The external force affects this route in two mutually opposing mechanisms: 1) The backward diffusion is accelerated by the external force as in eq. (5), which increases the transition to 2TDM0. 2) The MT-bound blue motor domain receives the half of external force in 2TDM0 state, but not in 2TDM+ state, which suggests that the 2TDM0 has extra energy cost due to external-force based factor *f*_*ext*_(*F* /2)=10^(*F pN*)^, which thus reduces the transition to 2TDM0. In fact, the two effects are nearly cancelled out, but, in our current estimate, the second effect is slightly larger. This inhibitory effect to 2TDM0 explains the slight increase in the transition frequency to 3T*DM+. We, however, note that our estimate in the second effect comes from the experimental data of detachment force for *Dictyostelium* dynein, which contain some uncertainty. If the detachment force were 3 pN, for example, instead of 2 pN used here, the second effect would be weaker than the first effect, and thus we may not see the slight increase in the velocity upon a weak external force.

### Effects of internal force and diffusion rates

Our model contains some parameters of which appropriate values are hardly known from experiments to date. Here, we address effects of two such parameter values; the internal force factor, *f* _∫¿¿_, and the diffusion rates, *λ*=*λ*_*for*_ =*λ*_*back*_ without the external force. As the default, we set *f* _∫ ¿=10000 ¿_ and *λ*=200. Here, we repeated the same simulations with three other parameter choices and compared them with the default one (S3 Fig). (1) *f* _∫ ¿=10000 ¿_ and *λ*=200 (the default set), (2) *f* _∫¿=1 ¿_ and *λ*=200, (3) *f* _∫ ¿=10000 ¿_ and *λ*=500, and (4) *f* _∫ ¿=10000 ¿_ and *λ*=1000. We plot the results for run-lengths (S3A and S3B Fig) and velocities (S3C and S3D Fig) as functions of [ATP] (S3A and S3C Fig) and the external force (S3B and S3D Fig). The [ATP] dependence was tested with no external force. The external force dependence was investigated with [ATP] = 1 mM.

At first, the two major features were kept for all the cases. 1) The model dynein does not move uni-directionally without ATP. 2) The run-length decreased as [ATP] increased.

Looking into the effect of the diffusion rate *λ*, we find that all the results for the cases (1), (3), and (4) are rather similar each other, suggesting that the diffusion rate value does not affect the overall behavior significantly.

When we removed the effect of internal force *f* _∫¿¿_ in (2)(green curves in S3 Fig), the run-length at higher [ATP] becomes longer, compared to the default set (1) that contains internal force (S3A Fig), while the velocity is significantly lower than the default case (S3C Fig). Thus, the system that lacks the internal force effect tends to bind more strongly to the MT, making the run-length longer and the velocity lower. The effect of the internal force, *f* _∫¿¿_, is to accelerate the dissociation of the lagging motor domain (*DM* →*D*), which facilitates the forward step of that domain increasing the velocity in one hand, and enhances the detachment of the entire dynein from MT thus decreasing the run length in the other hand.

Next, we examined the walking mechanisms in different *f* _∫¿¿_ and *λ* setups (S3E Fig). It is clear that in the system without the internal force *f* =1, the hand-over-hand moves become more prominent, while the inchworm moves are less probable, compared to the default case with the internal force. This is because the pathway; 1DMDM+ → 21DMD+ → 46DMD0 → 26DMDM0, observed frequently in the default case *f* =10000 is less prominent for the case of *f* =1, due to the deceleration of the first step in this pathway. In the transition 1DMDM+ → 21DMD+, the dissociation of the lagging motor domain is facilitated by the presence of the internal force. At the same time, when the lagging motor domain detaches from MT, it binds ATP leading to the hand-over-hand motions. Therefore, the internal force is important for the characteristic walking manner, inchworm motion.

Additionally, we also notice larger diffusion constant value shows slight increase in the inchworm motions. This is because the motor domain often moves diffusively twice within one ATP cycle in the case of large *λ* value, and so the lagging motor domain tends to detach via internal force and moves to the forward direction. When it then binds to MT, this pathway goes to 26DMDM0, regarded as the inchworm motions.

### Limitation and future prospect

As discussed, our kinetic model includes some parameters which are not derived from experiments or which are taken from experiments with mixed conditions such as different species or different ionic strengths. In particular, data on the effect of external force were missing for *Dictyostelium* dynein due to inherently weaker MT binding affinity of *Dictyostelium* dynein than those from yeast or human. However, since the time courses of movements have been measured, we could infer these model parameters via Bayesian approaches, which is kept for future studies.

In addition, for simplicity, we limit ourselves the relative positions of the two motor domains being +8 nm, 0 nm, or −8 nm. It is straightforward to extend it to 16 nm or 24 nm to make it closer to experimental estimates. Limiting the relative position to discrete states is perhaps not an ideal setup. Extending it to continuous value would be desired. Moreover, it has been known that dynein occasionally make sidesteps to different protofilaments, which, for simplicity, we have not included in the model.

## Conclusion

We proposed a kinetic model for bipedal motions of cytoplasmic dynein, simulated it via the Gillespie Monte Carlo method obtaining results consistent with many of previous motility experiments. The detailed pathway analysis provided new and versatile molecular mechanisms of bipedal motions of dynein, including the inchworm motions and the hand-over-hand motions. The kinetic model contains the 5 states of each motor domain based on the ATP-dependent structural changes in the linker and the MTBD, as well as 3 states for relative position of the two motor domain, resulting into 5 x 5 x 3 = 75 states together. The master equation in this 75 states was solved by the Gillespie algorithm. As a result, with a single parameter set, we successfully reproduced many characteristic behavior of dynein found in previous experiments; the inchworm motions as the dominant mode, the hand-over-hand motions, backsteps, and stagnation. Our model suggests that, in the prominent inchworm movement, only the leading motor domain moves via the ATP-dependent linker power-stroke motions, while the lagging motor domain is moved diffusively dragged by the leading domain. In addition, the hand-over-hand motion in the model dynein is distinct from that of kinesin by the usage of the power-stroke.

## Supporting Information

### Supporting Figure Caption

**Fig S1: Effects of internal forces.** The model takes into account the asymmetric detachment rates of the motor domain; when both domains are bound on MT, the lagging domain dissociates more rapidly than the leading domain. In the master equation approach, the effect is real-ized by introducing a multyping/dividing factor, *f*_int_ in the transition rates, which implies the changes in the free energy by ln (*f*_int_). Taking care of the detailed balance, we need to introduce the same factor in many transitions connected in the kinetic network. The introduced multyplying/dividing factors are defined in the figure.

Fig S2: All the 10000 trajectories superimposed for various [ATP] and external forces.

Fig S3: Motility with different model parameters are examined.

